# Coupled Mixed Model for Joint Genetic Analysis of Complex Disorders with Two Independently Collected Data Sets

**DOI:** 10.1101/336727

**Authors:** Haohan Wang, Fen Pei, Michael M. Vanyukov, Ivet Bahar, Wei Wu, Eric P. Xing

## Abstract

In the last decade, Genome-wide Association studies (GWASs) have contributed to decoding the human genome by uncovering many genetic variations associated with various diseases. Many follow-up investigations involve *joint analysis* of multiple independently generated GWAS data sets. While most of the computational approaches developed for joint analysis are based on summary statistics, the joint analysis based on individual-level data with consideration of confounding factors remains to be a challenge. In this study, we propose a method, called Coupled Mixed Model (CMM), that enables a joint GWAS analysis on two independently collected sets of GWAS data with different phenotypes. The CMM method does not require the data sets to have the same phenotypes as it aims to infer the unknown phenotypes using a set of multivariate sparse mixed models. Moreover, CMM addresses the confounding variables due to population stratication, family structures, and cryptic relatedness, as well as those arising during data collection such as batch effects that frequently appear in joint genetic studies. We evaluate the performance of CMM using simulation experiments. In real data analysis, we illustrate the utility of CMM by an application to evaluating common genetic associations for Alzheimers disease and substance use disorder using datasets independently collected for the two complex human disorders. Comparison of the results with those from previous experiments and analyses supports the utility of our method and provides new insights into the diseases.

The software is available at https://github.com/HaohanWang/CMM

## 1 Introduction

Genome-wide Association Studies (GWASs) have helped reveal about 10,000 associations between genetic variants in the human genome and diseases (Visscher *et al*., 2017). With the success of GWASs involving analysis of single data sets, a natural follow-up is to investigate multiple data sets (Wu *et al*., 2014), which we refer to as joint analysis. A joint analysis may uncover genetic mechanisms that cannot be discovered in a single analysis (Mukherjee *et al*., 2013). For example, recent studies have revealed overlapping genetic factors that influence multiple psychiatric disorders (Pain *et al*., 2018), genetic correlations between schizophrenia, ADHD, depression, and cannabis use (Walters *et al*., 2018), as well as an association between schizophrenia and illicit drug use (Mallard *et al*., 2018). Also, co-occurrences of substance use disorders (SUDs) and psychopathology have been observed in national epidemiologic surveys (Grant *et al*., 2015, 2016), which suggests a further joint analysis should be conducted to uncover potential common genetic factors underlying both SUDs and diseases involving cognitive dysfunction.

However, a joint genetic analysis using two independently collected data sets can be very challenging. In addition to the issues commonly expected from single data sets such as population stratification (Devlin and Roeder, 1999), a straightforward application of computational methods proposed for single data sets for the joint analysis could result in false discoveries caused by confounding factors such as the batch effects due to different data collection procedures. Moreover, two independently collected data sets do not often share the phenotypes of interest. To help better understand these challenges, we illustrate them in detail in Fig. 1. For the two data sets 1 and 2 originally collected for independent studies of the red and blue phenotype, respectively, a joint analysis aims to discover common genetic variants associated with both of these phenotypes. However, in order to perform such analysis, as shown in Fig. 1, all the information that is enclosed in the boxes with dashed lines needs to be inferred which could pose major challenges for these analyses; questions need to be answered involve, *e.g*., what is the blue phenotype of the samples in data set 1 since the blue phenotype may not be collected when the data set 1 is generated? How to deal with different confounding factors present in different data sets, including population stratification, family structures, cryptic relatedness, and data collection confounders?

**Figure 1:**
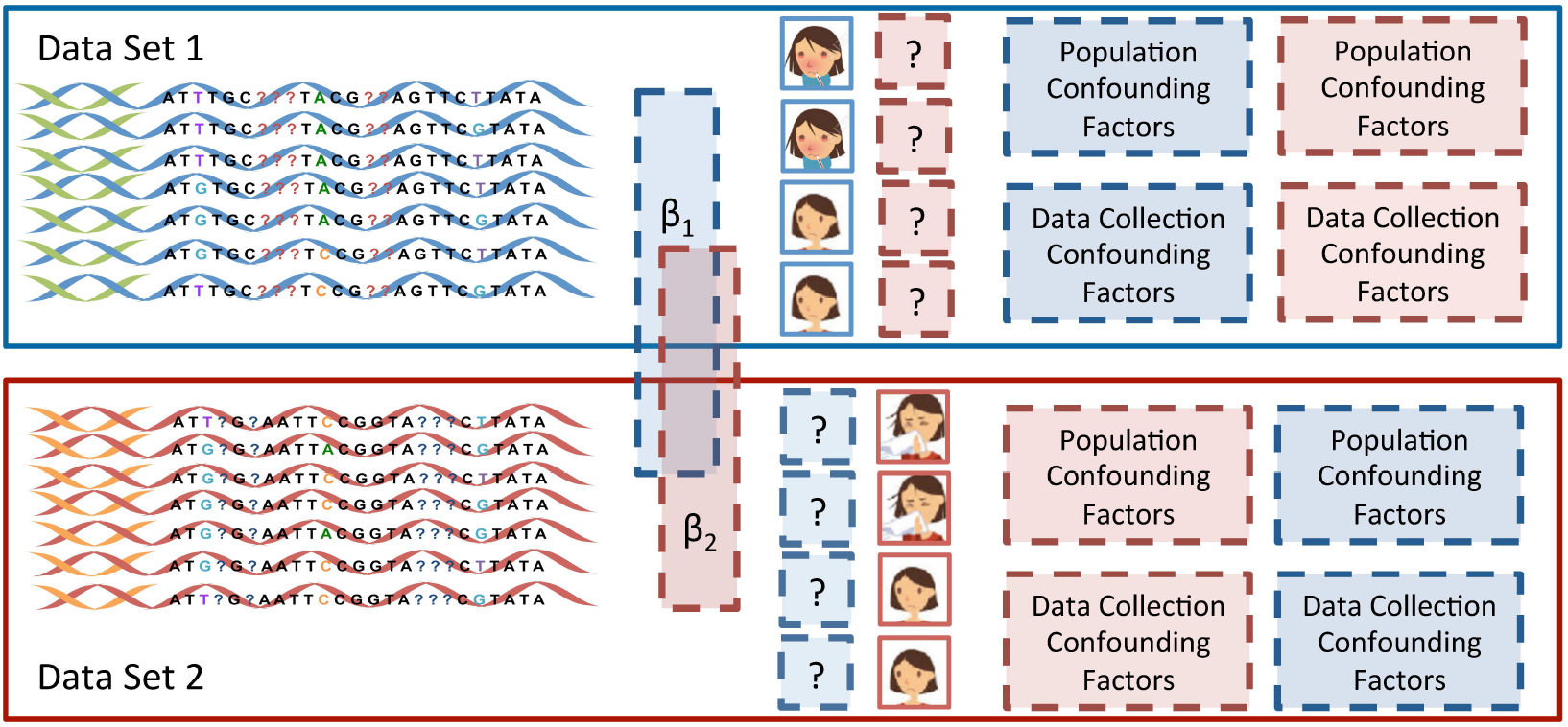
Illustration of the existing challenges when conducting a joint analysis on two independently collected data sets with two different phenotypes.

Existing methods for joint analysis on genetic data are mostly built on summary statistics (e.g., McGeachie *et al*., 2014; Giambartolomei *et al*., 2014; Kang *et al*., 2014; Zhu *et al*., 2015; Bulik-Sullivan *et al*., 2015; Nieuwboer *et al*., 2016; Hu *et al*., 2017; Wen *et al*., 2017; Liu *et al*., 2017; Sha *et al*., 2018; Guo and Wu, 2018) More recently, Turley *et al*. (2018) introduced multi-trait analysis of GWAS (MTAG) that can perform joint analysis using the summary statistics calculated from cohorts with overlapping samples. Zeng *et al*. (2018) proposed a regularized Gaussian mixture model called iMAP to infer the association between SNPs to correlated phenotypes. Qi and Chatterjee (2018) presented a heritability-informed power optimization method that finds an optimal linear combinations of association coefficients.

While summary statistics can help uncover common genetic factors from joint analysis, individual-level data nonetheless contains more information that allows the analyst to adjust for patient-level covariates, repeated measures, *etc*. (Siddique *et al*., 2015). Recently, Dai *et al*. (2018) proposed a method for joint analysis which integrates individual-level data together with summary-level data. Yang *et al*. (2018) directly used individual-level data for the joint analysis of traits that are collected separately from different cohorts. However, none of these methods took advantage of the rich information of the distribution of SNPs in the individual-level data, which allows the analyst to infer and correct the sample population structure or other potential confounding factors. In this work, we introduce a computational method for joint genetic analysis using individual-level data with correction of potential confounding factors.

Here, we propose a method, namely Coupled Mixed Model (CMM), for a joint association analysis that directly operates on two GWAS sequence data sets. CMM aims to address all the challenges above and to provide a reliable joint analysis of the data sets by inferring the missing information as illustrated in Fig. 1. In particular, CMM infers the missing phenotypes and various confounding factors with the maximum likelihood estimation. It is also noteworthy that our method is different from the approaches for missing phenotype imputation such as (Dahl *et al*., 2015; Hormozdiari *et al*., 2016) in that our method aims to address the challenges when there are no empirical data which allows the correlation between different phenotypes to be measured – a common situation for independently collected data sets which are not originally generated for joint analysis purposes. We first verify the performance of our methods with simulation experiments, and then apply our method to real GWAS data sets previously generated for investigating genetic variants associated with substance use disorders (SUDs) and Alzheimer’s disease (AD), respectively, for joint analysis.

## 2 Materials and methods

### 2.1 Coupled Mixed Model

The following are the notations we use in this work: The *subscript* denotes the *identifier of data set*, and the *superscript in parentheses* denotes the *identifier of phenotypes*. Genotypes and phenotypes are denoted as **X** and **y**, respectively. Also, *n* denotes the sample size, and *p* denotes the number of SNPs. Specifically, consider a scenario as illustrated in Fig. 1, **X**_1_ and **X**_2_ represent the genotypes of the samples in data sets 1 and 2 with the dimension of *n*_1_ × *p* and *n*_2_ × *p*, respectively. 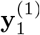 and 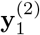 denote the vectors of phenotypes 1 and 2, respectively, of the dimension *n*_1_ × 1 for the samples in data set 1. Note that 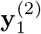 is not observed. Similarly, 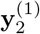 and 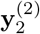 denote the vectors of phenotypes 1 and 2, respectively, of the dimension *n*_2_ × 1 for the samples in data set 2. 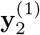 is not observed.

Our method does not require *n*_1_ = *n*_2_. However, for the convenience of the discussion, we will assume *n*_1_ = *n*_2_ = *n*. The case of *n*_1_ ≠ *n*_2_ can be easily generalized by weighing the corresponding cost function components with 1/*n*_1_ and 1/*n*_2_, respectively. Following the similar logic, we introduce our method with the simplest linear models, but our method can be extended to the case of generalized linear models; for example, for case-control data, one can directly apply our method to binary trait data, as done by many previous examples (Moser *et al*., 2015; Speed and Balding, 2014; Weissbrod *et al*., 2016; Zhou *et al*., 2013; Zeng *et al*., 2018). Also, one can use our method with the residual phenotype after regressing other additional covariates (*e.g*, age or sex).

Straightforwardly, for the scenario shown in Fig. 1, we have:

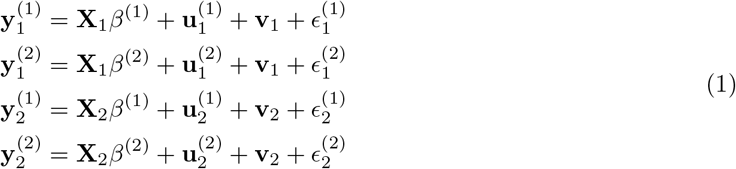

where 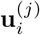 accounts for the confounding effects due to population stratification, family structures and cryptic relatedness in data set *i* with phenotype *j*; and **v**_*i*_ accounts for the confounding effects due to data collection (*e.g*., batch effects) in data set *i*; 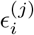 stands for residual noises for data set *i* with phenotype *j*, and 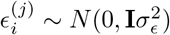, where **I** is an identity matrix with the shape of *n* × *n*. Notice that we will drop the unidentifiable term 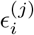 later during parameter estimation, otherwise these terms will turn the entire model unidentifiable.

We have 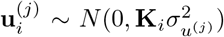 for data set *i* with phenotype *j*. As observed by Devlin and Roeder (1999), population stratification can cause false discoveries because there exist real associations between a phenotype and untyped SNPs that have similar allele frequencies with some typed SNPs that are not actually associated with the phenotype, which, as a result, can lead to false associations between the phenotype and the typed SNPs. Since these false associations due to confounders from population stratification are phenotype specific, we model 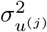 as phenotype-specific. Hence, although we have four different variance terms (*i.e*., 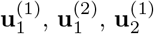 and 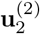) accounting for population confounders, they are only parameterized by two scalars, 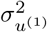 and 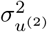. 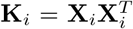 is the kinship matrix, constructed following the genetics convention (Yang *et al*., 2014). A more sophisticated construction of the kinship matrix may be used to improve detection of the signals, but these details are beyond the scope of this paper. One can refer to examples in (Listgarten *et al*., 2013; Tucker *et al*., 2014; Wang *et al*., 2017) for more details.

To model the confounders due to data collection, we have 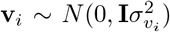 for data set *i*. Because these confounders are only related to the data collection procedure, we model 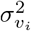 as data set-specific.

For the independently collected data sets, we only observe 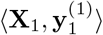 and 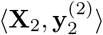. Since we are interested in estimating *β*^(1)^ and *β*^(2)^, we also need to estimate 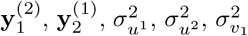, and 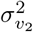 Eq. 1. As noted earlier, we drop the 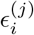. to avoid the model to become unidentifiable.

In order to estimate *β*^(1)^ and *β*^(2)^, we minimize the joint negative log-likelihood function. Because there are only a subset of SNPs that contribute to the phenotype, we introduce the standard *ℓ*_1_ regularization by setting the prior distribution of *β*^(1)^ and *β*^(2)^ as a Laplace distribution. Additionally, to encourage our method to find common SNPs associated with both phenotypes, we use a simple constraint, as shown in Constraint (3). Taken together, the optimization problem for solving our model in Eq. 1 can be represented as follows:

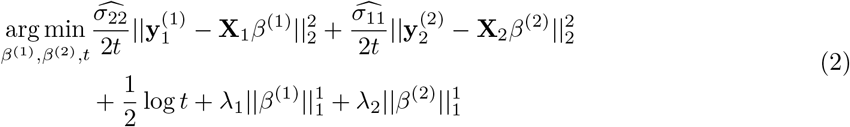

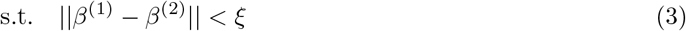

where

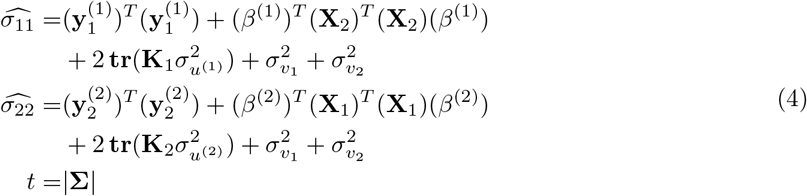

where Σ is the covariance matrix defined as:

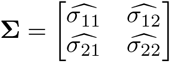

and we have:

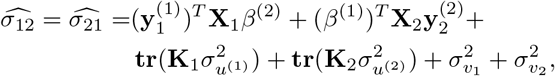

and *ξ* denotes a small number. The detailed derivation is described in Supplement Section S1. The key steps involve replacing 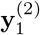 with **X**_1_*β*^(2)^, and replacing 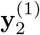 with **X**_2_*β*^(1)^, and then writing out the joint likelihood function of Equation 1.

To solve the optimization Function (2), we propose a strategy as follows. We first estimate the parameters 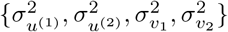 following the P3D set-up (Zhang *et al*., 2010). Then we propose an iterative updating algorithm that decouples the dependency between {*β*^(1)^, *β*^(2)^} and *t* in the optimization function 2 and solves for {*β*^(1)^, *β*^(2)^} and *t* with ADMM (Boyd *et al*., 2011), which naturally uses the Constraint 3. We also offer a proof to show that our iterative updating algorithm will converge. The details of the algorithm and the convergence proof are presented in the Supplement Section S2 and S3, respectively.

### 2.2 Implementation

The implementation of the CMM method is available as a python software. Without installation, one can run the software with a single command line. It takes binary data in a standard Plink format for each of the two data sets as input. If there are mismatched SNPs between the data sets, CMM will use the intersection of these SNPs. We recommend the users to query CMM to identify a specific number of SNPs for each data set and CMM can tune the hyperparameters accordingly (Wang *et al*., 2018). However, users can also choose to specify the regularization parameters. If none of the above information is specified, CMM will automatically conduct five-fold cross-validation to tune parameters. *ξ* does not need to be specified or tuned, because it can be dropped due to ADMM. The implementation is available as a standalone software^1^. More detailed instructions of how to use the software are presented in Supplement Section S4.

In theory, the computational complexity of the first step of the algorithm is *O*(*n*^3^), and complexity of the second step is *O*(*np*). In practice, as we observe on two data sets with hundreds of samples and 200k SNPs, it takes CMM around a minute to converge given a set of hyperparameters on a modern server (2.30GHz CPU and 128G RAM, Linux OS), and up to an hour to finish the entire hyperparameter tuning process.

## 3 Results

### 3.1 Simulation Experiments

We compare CMM to several approaches using simulated data sets.

- HG(W): Joint analysis conducted with the hypergeometric tests (McGeachie *et al*., 2014) when the two independent problems are each solved by the standard univariate Wald testing with the Benjamini-Hochberg (BH) procedure (Benjamini and Hochberg, 1995). This is the most popular approach in GWAS for a single data set.
- HG(L): Joint analysis conducted with the hypergeometric tests (McGeachie *et al*., 2014) when the two independent problems are each solved by a standard linear mixed model with the Benjamini-Hochberg (BH) procedure (Benjamini and Hochberg, 1995).
- CD: Combining data-set approach. CD merges two data sets *X*_1_ and *X*_2_ into one *X* = [*X*_1_; *X*_2_] and create a pseudo phenotype *y* ∈ {0, 1}^*n*_1_+*n*_2_^ where *y*(*i*) = 1 if the *i*^th^ sample has either one of the two diseases.
- iMAP: integrative MApping of Pleiotropic association, which is a method for joint analysis that models summary statistics from GWAS results by integrating SNP annotations in the model (Zeng *et al*., 2018). For a fair comparison of the methods, we do not use the SNP annotations with this method.
- MTAG: multi-trait analysis of GWAS (Turley *et al*., 2018), which is also a method for joint analysis of GWAS data sets using summary statistics, which accounts for potential confounders due to population stratification or cryptic relatedness.
- LR: *ℓ*_1_-regularized logistic regression, which can be directly applied to the two independent data sets for joint analysis. We select the intersection of the identified SNPs associated with each of the phenotypes as the SNPs jointly associated with both phenotypes.
- AL: Adaptive Lasso, which is an extension of the Lasso that weighs the regularization term (Zou, 2006) (enabled by the method introduced in (Huang *et al*., 2008) for high-dimensional data). AL is applied to the independent data sets in the same manner as LR. We use the logistic-regression version of the method if the phenotypes are binary.
- PL: Precision Lasso, a novel variant of the Lasso, that is developed for analyzing data with correlated and linearly dependent features, commonly seen in genomic studies (Wang *et al*., 2018). PL is applied to the independent data sets in the same manner as LR.
- JL: Joint Lasso, which is a method we implement in this study for a fair comparison of our proposed CMM method. JL solves the lasso problems jointly with the constraint *β*^(1)^ = *β*^(2)^ with ADMM. This approach can be seen as a CMM method without consideration of the confounding factors in the data.
- CMM: Coupled Mixed Model. Our proposed method.

We simulate two independent data sets with binary phenotypes, whose SNPs are generated via SimuPop (Peng and Kimmel, 2005) with population structures. We also introduce the influences from confounding factors, resulting in a roughly 0.25 signal-to-noise ratio for effect sizes. We mainly experiment with two different settings: the number of the associated SNPs and the fraction of these SNPs that are jointly associated with both phenotypes. We repeat the experiments with 10 different random seeds. Details of simulation are in Supplement Section S5.1.

We first evaluate these methods with the focus on finding the SNPs associated with both phenotypes, and compare the performance of the competing methods with ROC curves. For the univariate testing methods (HG, CD, MTAG), the curves are plotted by varying the null-hypothesis-rejecting threshold of p-values, while for multivariate regularized regression methods, the curves are plotted by varying the regularizing hyperparameter (200 different choices evenly distributed in logspace from 10^−5^ to 10^5^).

Fig. 2 shows the ROC curves of the compared methods in terms of their abilities to find the SNPs associated with both phenotypes. Overall, the results favor our CMM method significantly. In comparison with the other methods, the superiority of the proposed CMM is more evident when there are fewer associated SNPs in each data set, and also when there are fewer SNPs associated with both phenotypes. For example, as shown in Fig. 2, when only 0.1% of the SNPs are associated with a phenotype (first row), the advantage of CMM can be clearly seen; however, when 1% of SNPs are associated with a phenotype (last row), CMM barely outperforms HG(L) methods.

**Figure 2:**
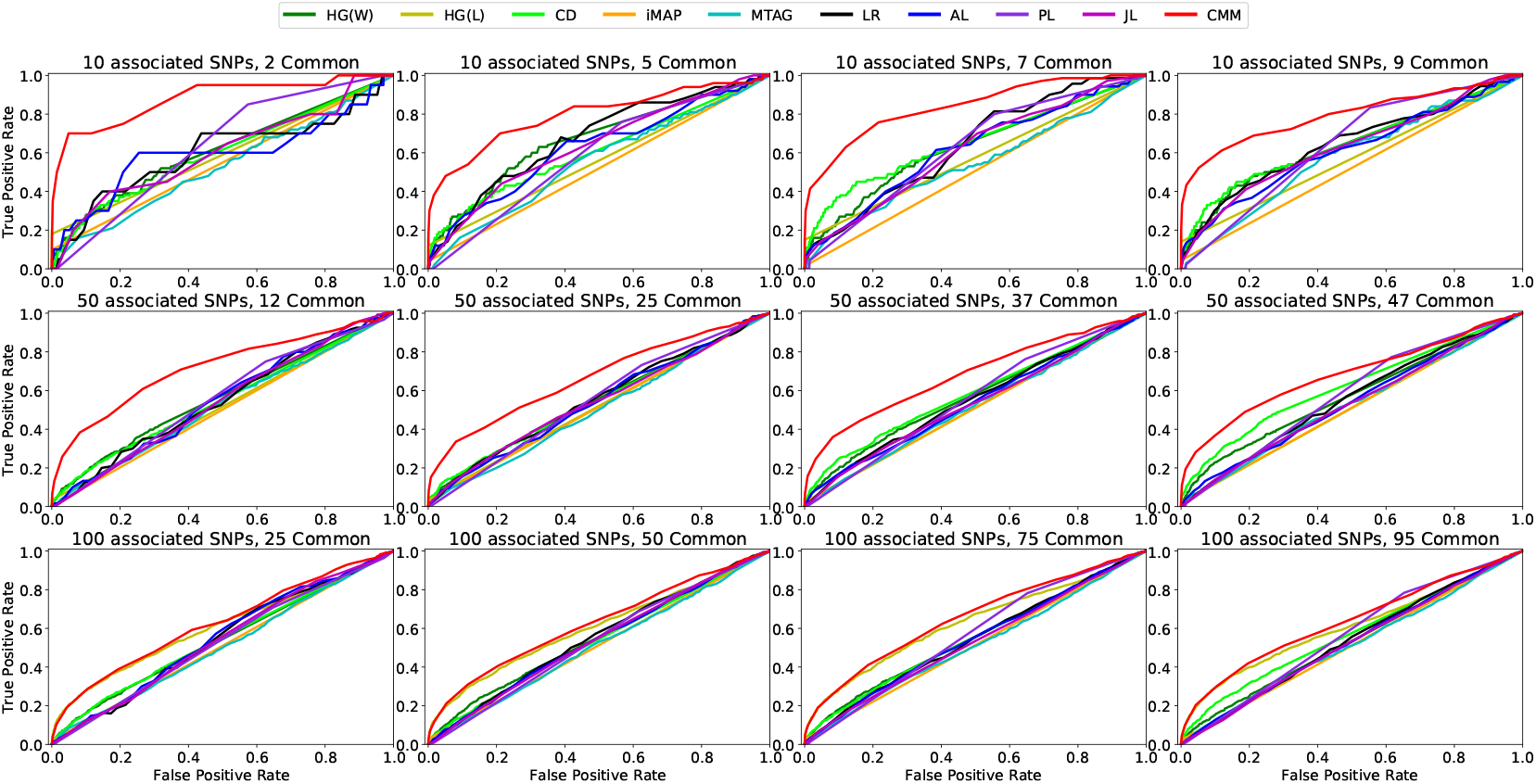
The ROC curves of the compared methods in terms of identifying the SNPs that are jointly associated with both phenotypes.

By comparing the performances of the compared methods in different columns in Fig. 2, we can see how the common SNPs (*i.e*., those associated with both phenotypes) affect the results: as the percentage of the common SNPs increases, in general, the performances of all the compared methods increase. Also, we notice that the performance of CMM does not vary significantly as the number of the common SNPs varies, This observation indicates that the Constraint (3) in our optimization problem does not necessarily deteriorate the method’s performance even when the two phenotypes are less related.

With the clear advantage of CMM, we now proceed to discuss more about the other competing methods. We notice that multivariate methods (LR, AL, JL) tend to perform well when there are less associated SNPs as well as less common SNPs, while univariate methods (HG(W), HG(L), CD) favor the opposite scenarios with more associated SNPs and more common SNPs. For instance, JL, which can be considered as a multivariate version of CD, barely outperforms CD. As the number of the common SNPs increases, the performance of CD improves clearly, while that of JL does not. This result can be explained as follows: CD only aims to recover the common SNPs, while JL balances between minimizing the two logistic regression cost functions and minimizing the differentiation between coefficients which may not result in a more effective recovery of the common SNPs. Unfortunately, summary statistics-based methods (iMAP and MTAG) do not perform well in our simulation experiment settings, most likely due to the presence of the multiple sources of confounding factors in the simulated data. Also, iMAP is introduced as a method which leverages the power of SNP annotations for joint analysis, but we do not include the annotation information in the experiments for fair comparisons.

We also notice that LMM performs surprisingly well when there are many associated SNPs. For example, when there are 1% of the associated SNPs (last row of Fig. 2), LMM performs as the second best method. However, LMM does not perform well with fewer associated SNPs, as shown in the first two rows of Fig. 2. Furthermore, we plot the results of the ROC curves of the compared methods regarding their abilities in uncovering the associated SNPs separately for each data set, which are shown in Supplement Section S4.2. Together, these simulation results demonstrate that CMM outperforms the other methods in terms of finding common SNPs associated with both phenotypes, as well as finding associated SNPs with individual phenotype.

We also tested our CMM method for predicting the phenotypes across data sets in comparison to the other competing regression-based methods. The results are presented in Supplement Section S5.2 and S5.3.

### 3.2 Real Data Analysis: Joint Genetic Analysis for Alzheimer’s disease and substance use disorder

#### 3.2.1 Application of CMM to two GWAS data sets for AD and SUDs

In the real data analysis, we apply our proposed CMM method to two GWAS data sets independently generated previously to investigate genetic association for AD and SUDs, respectively. The AD data set was collected from the Alzheimers Disease Neuroimaging Initiative (ADNI)^2^ and the SUD data set was collected by the CEDAR Center at the University of Pittsburgh^3^. For the AD data set, we only used the data generated from the individuals diagnosed with either AD or normal controls. There are 477 individuals in the final AD data set with 188 case samples and 289 control samples. For the SUD data set, we consider the subjects with drug abuse history as the case group and the subjects with neither drug abuse nor alcohol abuse behavior as the control group, excluding the subjects with only alcohol abuse behavior (but not drug abuse history), because alcoholism is usually believed to be related to drug abuse. There are 359 patients in the final SUD data set with 153 case samples and 206 control samples. We also exclude the SNPs on X-chromosome following suggestions of previous studies (Bertram *et al*., 2008). There are 257361 SNPs in these two data sets left to be examined. Even though the sample sizes of the AD and the SUD data sets are small, which unfortunately is a common situation for genetic studies of complex human diseases, particularly for SUDs, our results suggest that our CMM method can help identify promising genetic variants that are worth further investigation.

Due to the statistical limitation of selecting hyperparameters using cross-validation and information criteria in high dimensional data (Wang *et al*., 2018), we tune the hyperparameters according to the number of SNPs we aim to select, following previous work (Wu *et al*., 2009; Marchetti-Bowick *et al*., 2016; Wang *et al*., 2018) and the hyperparameters of our model will be tuned automatically through binary search for the set of parameters according to the number of SNPs we inquire. This hyperparameter selection procedure has been shown to generate less false positives in general than cross-validation, even when the queried number of SNPs is (reasonably) misspecified (Wang *et al*., 2018). To mitigate the computation load, the algorithm will terminate the hyperparameter search when the number of the reported SNPs lies within 50% to 200% of the number we inquire.

We inquire for 30 SNPs selected in each data set, and CMM identified five SNPs that are associated with both SUD and AD, which is reported in Table 1. CMM reported 15 additional SNPs and 35 additional SNPs for SUD and AD respectively, which are reported in Tables S1 and S2 (Supplement Section S7). Notably, we do not find much overlap between our findings and those from the previous studies in the GWAS Catalog (Welter *et al*., 2013), and we believe this is because our method explicitly favors to identify the SNPs that are jointly associated with both of the disease phenotypes. Nevertheless, we find many pieces of evidence supporting our findings. The following discussion focuses on the validation of these five identified SNPs.

**Table 1:** The SNPs that the CMM method identifies from both the SUD and the AD data sets. The SNPs are ranked by the absolute values of their estimated effect sizes, and showed in the “SUD rank” and “AD rank” columns. The information of whether a SNP is located within a region of a gene is taken from the Database for Single Nucleotide Polymorphisms (dbSNP) (Sherry *et al*., 2001), and listed in the “Gene” column.

| SNP | SUD rank | AD rank | Chr. | Chr. Position | Gene |
| --- | --- | --- | --- | --- | --- |
| rs2131691 | 1 | 1 | 11 | 26574855 | *ANO3* |
| rs1709317 | 5 | 8 | 2 | 23536638 | *KLHL29* |
| rs4713797 | 6 | 10 | 6 | 34490756 | *PACSIN1* |
| rs224534 | 12 | 3 | 17 | 3583408 | *TRPV1* |
| rs1057744 | 16 | 11 | 14 | 105150705 | *JAG2* |

#### 3.2.2 Validation of the identified common SNPs associated with both AD and SUDs

##### Statistical validation

In order to validate the five identified common SNPs, we first compared the distribution differences of SNPs between the case and control samples in each of the diseases. We notice that in most cases, the allele frequencies are different between the case and the control samples (shown in Table 2). Also, we examine the statistical significance of independence between the SNPs in the control group vs. the case group with the student’s t-test. Seven out of the ten tests report a statistically significant sign of independence (shown in Table 2).

**Table 2:** The minor allel frequencies (MAFs) of the five identified SNPs in the case (“AD” column) and the control (“C” column) samples. The overall MAFs (in “all” column are reported for reference. The p-values of the student’s t-tests are also reported. The statistically significant p-values which are below the threshold of 0.05 are shown in bold.

|  | AD |  |  |  | SUD |  |  |  |
| --- | --- | --- | --- | --- | --- | --- | --- | --- |
|  | all | C | AD | P-value | all | C | SUD | P-value |
| rs2131691 | 0.47 | 0.41 | 0.44 | **4.78E05** | 0.47 | 0.46 | 0.48 | 4.64E-01 |
| rs1709317 | 0.36 | 0.31 | 0.45 | **6.82E-06** | 0.27 | 0.30 | 0.22 | **2.28E-02** |
| rs4713797 | 0.48 | 0.47 | 0.41 | **6.04E-04** | 0.41 | 0.45 | 0.37 | 5.32E-02 |
| rs224534 | 0.33 | 0.28 | 0.43 | **1.41E-06** | 0.25 | 0.26 | 0.25 | 7.91E-01 |
| rs1057744 | 0.49 | 0.45 | 0.42 | **4.81E-04** | 0.39 | 0.42 | 0.35 | **3.71E-02** |

##### Literature support

Due to the lack of direct information on the SNPs and the disease phenotypes, we also verify our findings via literature search based on the relationship between the genes where the identified SNPs reside and the phenotypes.

Our results show that *rs224534* identified by CMM to be associated with both AD and SUD resides in *TRPV1* which encodes transient receptor potential cation channel subfamily V member 1. Previous evidence showed that positive modulation of the *TRPV1* channels could be a potential target for mitigation of AD (Jayant *et al*., 2016), suggesting an important involvement of *TRPV1* in AD. In addition, Nguyen *et al*. (2014) have also shown that *TRPV1* plays a key role in morphine addiction. Blednov and Harris (2009) showed that the deletion of *TRPV1* in mice altered behavioral effects of ethanol which indicates a connection between *TRPV1* and alcoholism.

Moreover, *TRPV1* mediates long-term synaptic depression in the hippocampus (Gibson *et al*., 2008), which is key to reward-related learning and addiction (Kauer and Malenka, 2007). Further, we notice that in the “Inflammatory mediator regulation of TRP channels” pathway of the KEGG database (Kanehisa *et al*., 2016), *TRPV1* serves as a Ca2+ channel. Ca2+ binding to calmodulin (CaM) activates Ca2+/CaM-dependent protein kinase II (CAMKII). CaMKII is involved in many signaling cascades and is an important mediator of learning and memory (Yamauchi, 2005), which plays an important role in neuropsychiatric disorders including drug addiction, schizophrenia, depression and multiple neurodevelopmental disorders (Robison, 2014; Müller *et al*., 2016).

##### Additional evidence using an independent approach

In addition to the statistical and literature support, we also validate *TRPV1* as a SUD-related protein using an independent study of the drug-target interaction analysis.

In this drug-target interaction analysis, we identified the known ligands of the corresponding proteins of each gene through drug/ligand-target interactions compiled in DrugBank (Wishart *et al*., 2017) and STITCH (Szklarczyk *et al*., 2015) databases. In addition, predicted ligands with high confidence were obtained by applying a probabilistic matrix factorization (PMF) model (Cobanoglu *et al*., 2013) on known drug/ligand-target interactions in DrugBank and STITCH. The data and the method are accessible on our online server^4^. Among the identified known and predicted ligands, we focused on the drugs that are associated with either SUD or AD. The results show that 4 SUD-related drugs are known to interact with *TRPV1* and 5 SUD-related drugs are predicted to interact with *TRPV1*, which supports the association between *TRPV1* and SUD.

In particular, as illustrated in Fig. 3, our analysis shows that *TRPV1* is the known target of medical cannabis (plant use of marijuana), as well as three cannabinoids (nabiximols, cannabidivarin, and cannabidiol) in cannabis, according to the annotations in DrugBank. In the PMF prediction model, *TRPV1* is the predicted target of two cannabinoids (tetrahydrocannabivarin, cannabichromene) extracted from cannabis, two synthetic cannabinoids (dronabinol and nabilone) of Δ9-THC (another cannabinoid from cannabis), as well as a central nervous system (CNS) depressant (flunitrazepam). These drugs are commonly known as drugs of abuse, and thus these results help verify the association between *TRPV1* and SUD.

**Figure 3:**
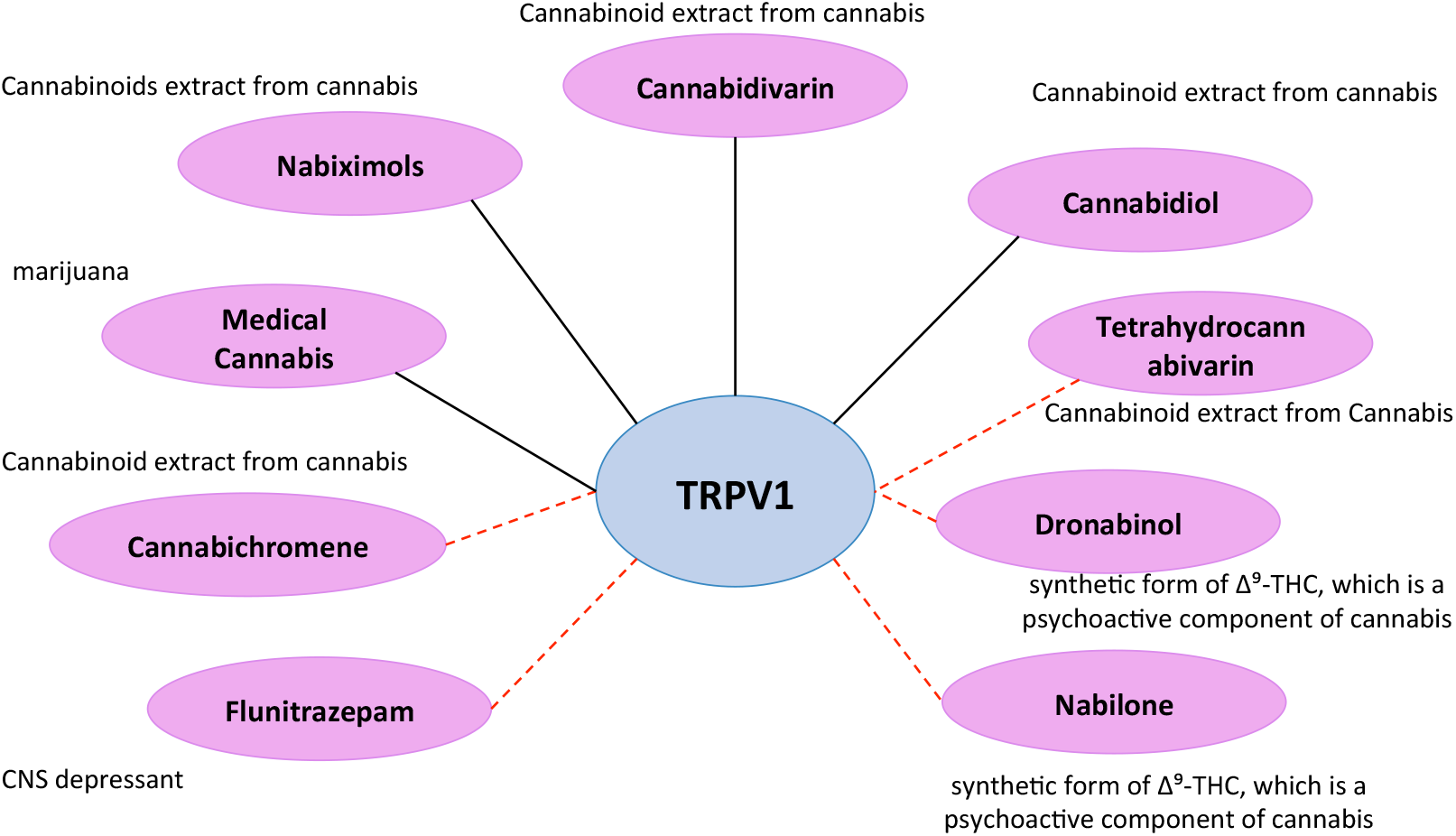
The interactions between *TRPV1* and 9 SUD-related drugs. Violet ellipses represent drugs of abuse; black solid edges represent known interactions in DrugBank; and red dashed edges represent predicted interactions using the PMF model.

Together, these results suggest that our findings, although explorative, may reveal novel genetic connections between SUD and AD. More discussions on the SNPs shown in Table 1 are presented in Supplement Section S6. For other SNPs identified by CMM which are associated with either AD or SUD, we discuss them in detail in Supplement Section S7.

## 4 Discussion and Conclusion

Following previous successes in joint genetic analysis using summary statistics-based approaches, we propose a novel method, Coupled Mixed Model (CMM), that operates on individual-level SNP data and aims to address challenges illustrated in Fig. 1. We further present an algorithm that allows an efficient parameter estimation of the objective function derived from our model.

With extensive simulation experiments, we showed the superior performance of the CMM method in comparison with several competing approaches. In the real data analysis, we applied our method to identify the common SNPs associated with both AD and SUD. CMM identified five SNPs associated with both of the disease phenotypes. Notably, one of the identified SNPs reside in the gene *TRPV1*, which has been linked to both AD and SUD by multiple pieces of evidence, including statistical tests showing differences in the allele frequencies between the case and the control samples, previous evidence in the literature, as well as results from an independent study of the drug-target interaction analysis. Together, we show that our proposed CMM method is able to uncover promising genetic variants that are associated with different disease phenotypes using individually collected GWAS data sets and reveal novel connections between diseases.

## Acknowledgements

The authors would like to thank Miaofeng Liu for help in the convergence proof of the algorithm. The authors would also like to thank Seunghak Lee and Ben Lengerich for suggestions and comments in the preparation of this manuscript. Data collection and sharing for this project was funded by the Alzheimer’s Disease Neuroimaging Initiative (ADNI) (National Institutes of Health Grant U01 AG024904) and DOD ADNI (Department of Defense award number W81XWH-12-2-0012). ADNI is funded by the National Institute on Aging, the National Institute of Biomedical Imaging and Bioengineering, and through generous contributions from the following: AbbVie, Alzheimers Association; Alzheimers Drug Discovery Foundation; Araclon Biotech; BioClinica, Inc.; Biogen; Bristol-Myers Squibb Company; CereSpir, Inc.; Cogstate; Eisai Inc.; Elan Pharmaceuticals, Inc.; Eli Lilly and Company; EuroImmun; F. Hoffmann-La Roche Ltd and its affiliated company Genentech, Inc.; Fujirebio; GE Healthcare; IXICO Ltd.; Janssen Alzheimer Immunotherapy Research & Development, LLC.; Johnson & Johnson Pharmaceutical Research & Development LLC.; Lumosity; Lundbeck; Merck & Co., Inc.; Meso Scale Diagnostics, LLC.; NeuroRx Research; Neurotrack Technologies; Novartis Pharmaceuticals Corporation; Pfizer Inc.; Piramal Imaging; Servier; Takeda Pharmaceutical Company; and Transition Therapeutics. The Canadian Institutes of Health Research is providing funds to support ADNI clinical sites in Canada. Private sector contributions are facilitated by the Foundation for the National Institutes of Health (www.fnih.org). The grantee organization is the Northern California Institute for Research and Education, and the study is coordinated by the Alzheimers Therapeutic Research Institute at the University of Southern California. ADNI data are disseminated by the Laboratory for Neuro Imaging at the University of Southern California.

## Footnotes

1 https://github.com/HaohanWang/CMM

2 http://adni.loni.usc.edu/

3 http://www.pitt.edu/cedar/

4 http://quartata.csb.pitt.edu

