## Supplement for "Coupled Mixed Model for Joint Genetic Analysis of Complex Disorders with Two Independently Collected Data Sets"

Supporting Information for *Joint Genetic Analysis of Complex Disorders from Two Independently Collected Data Sets*

#### S1 Derivation of Cost Function

In this section, we offer a detailed introduction of how CMM is derived. We continue with Equation 1 from the main manuscript, we write our the distribution of the data

$$\begin{aligned}
y_1^{(1)} &\sim N(X_1\beta^{(1)}, K_1\sigma^2_{u^{(1)}} + I\sigma^2_{v_1} + I\sigma^2_{\epsilon_1^{(1)}})\\
y_1^{(2)} &\sim N(X_1\beta^{(2)}, K_1\sigma^2_{u^{(2)}} + I\sigma^2_{v_1} + I\sigma^2_{\epsilon_1^{(2)}})\\
y_2^{(1)} &\sim N(X_2\beta^{(1)}, K_2\sigma^2_{u^{(1)}} + I\sigma^2_{v_2} + I\sigma^2_{\epsilon_2^{(1)}})\\
y_2^{(2)} &\sim N(X_2\beta^{(2)}, K_2\sigma^2_{u^{(2)}} + I\sigma^2_{v_2} + I\sigma^2_{\epsilon_2^{(2)}}),
\end{aligned}$$

and we introduce the following notations:

$$Y^{(1)} = \begin{bmatrix} y_1^{(1)} \\ y_2^{(1)} \end{bmatrix} \quad Y^{(2)} = \begin{bmatrix} y_1^{(2)} \\ y_2^{(2)} \end{bmatrix}$$

where we discard the residue term $\epsilon$.

Further, we can write out the joint likelihood function of $Y^(1)$ and $Y^(2)$, as following:

$$l(Y^{(1)}, Y^{(2)}, \beta^{(1)}, \beta^{(2)}) = -\frac{1}{2}\begin{bmatrix} Y^{(1)} - X\beta^{(1)} \\ Y^{(2)} - X\beta^{(2)} \end{bmatrix}^T \Sigma^{-1} \begin{bmatrix} Y^{(1)} - X\beta^{(1)} \\ Y^{(2)} - X\beta^{(2)} \end{bmatrix} - \frac{1}{2}\log|\Sigma| + c$$

where $X = [X_1, X_2]^T$, $c$ is a constant, and $\Sigma$ is defined as in Equation 2 in the main manuscript.

Further, since we have

$$\Sigma^{-1} = \frac{1}{|\Sigma|}\begin{bmatrix} \widehat{\sigma_{22}} & -\widehat{\sigma_{12}} \\ -\widehat{\sigma_{12}} & \widehat{\sigma_{11}} \end{bmatrix},$$

where $\sigma_{11}$, $\sigma_{12}$, $\sigma_{22}$ are defined in Equation 2 in the main manuscript.

With linear algebra, we will have:

$$\begin{aligned}
l(Y^{(1)}, Y^{(2)}, \beta^{(1)}, \beta^{(2)}) = &-\frac{1}{2t}\Big(\widehat{\sigma_{22}}(Y^{(1)} - X\beta^{(1)})^T(Y^{(1)} - X\beta^{(1)}) - \widehat{\sigma_{11}}(Y^{(2)} - X\beta^{(2)})^T(Y^{(2)} - X\beta^{(2)})\\
&- 2\widehat{\sigma_{12}}(Y^{(1)} - X\beta^{(1)})^T(Y^{(2)} - X\beta^{(2)})\Big) - \frac{1}{2}\log t + c
\end{aligned}$$

where $t = |\Sigma|$. Then we replace $y_2^{(1)}$ and $y_1^{(2)}$ with $X_2\beta^{(1)}$ and $X_1\beta^{(2)}$ respectively and drop the constant term and get:

$$l(y_1^{(1)}, y_2^{(2)}, \beta^{(1)}, \beta^{(2)}) = \frac{\widehat{\sigma_{22}}}{2t}||y_1^{(1)} - X_1\beta^{(1)}||_2^2 + \frac{\widehat{\sigma_{11}}}{2t}||y_2^{(2)} - X_2\beta^{(2)}||_2^2 + \frac{1}{2}\log t.$$

Finally, we introduce the model assumptions of the effect sizes, which constrains $\beta^{(1)}$ and $\beta^{(2)}$ to be sparse, and requires $\beta^{(1)}$ and $\beta^{(2)}$ to have common support, which leads to the Equation 2 introduced in the main manuscript.

### S2 Algorithm

#### S2.1 Estimating variances of the population confounders and data collection confounders

Despite the complicated cost function, solving for

$$\{\sigma^2_{u^{(1)}}, \sigma^2_{u^{(2)}}, \sigma^2_{v_1}, \sigma^2_{v_2}\}$$

is relatively straightforward because we have:

$$y_1^{(1)} = X_1\beta^{(1)} + u_1^{(1)} + v_1$$
$$y_2^{(2)} = X_2\beta^{(2)} + u_2^{(2)} + v_2$$

where $y_1^{(1)}$, $X_1$, $y_2^{(2)}$, and $X_2$ are all fully observed.

Equivalently, we have:

$$y_1^{(1)} \sim N(X_1\beta^{(1)}, K_1\sigma^2_{u^{(1)}} + I\sigma^2_{v_1})$$
$$y_2^{(2)} \sim N(X_2\beta^{(2)}, K_2\sigma^2_{u^{(2)}} + I\sigma^2_{v_2})$$

Thus, we can follow the conventional assumption that assumes fixed effects to be zero [1, 2, 3] for solving linear mixed models to estimate $\{\sigma^2_{u^{(1)}}, \sigma^2_{u^{(2)}}, \sigma^2_{v_1}, \sigma^2_{v_2}\}$. $K_i$ is usually constructed from $X_iX_i^T$ in genomic applications [4], which may introduce an "over-representing problem" since $X_iX_i^T$ has a full rank and hence represents the relationship between every pair of the samples in the data set [5]. Therefore, we use a truncated-rank approach proposed in [6] to reduce the rank of $K$. Specifically, we set the non-dominant eigenvalues of $K$ to be zero with a simple inspection of the slope of the eigenvalues as follows: if $\frac{S_i - S_{i+1}}{S_0} \leq \frac{1}{n}$, where $n$ is the number of samples, then we set the eigenvalue $S_i$ to be zero.

**An iterative updating algorithm for estimating effect sizes and the covariance matrix of the joint likelihood** We only have $\{\beta^{(1)}, \beta^{(2)}, t\}$ left to be estimated, however, directly estimating $t$ is difficult since it involves in four coupled terms in Function **??**. To simplify the problem, we introduce an approximation to decouple the dependencies among $t$, $\beta_i$ and $\sigma_{ii}$, leading to a neat solution involving two steps that can be conducted iteratively until convergence. The proof of the convergence of the algorithm is in Supplement Section S2.

**Calculating $t$ and $\sigma_{ii}$ given $\beta_i$** Calculating $t$ and $\sigma_{ii}$ is straightforward because both calculations can be done analytically. The analytical form for $\sigma_{ii}$ is shown in Eq. **??**. The analytical form of $t$ can be derived by equating the derivative of this following convex function to zero:

$$\frac{\widehat{\sigma_{22}}}{2t}||y_1^{(1)} - X_1\beta^{(1)}||_2^2 + \frac{\widehat{\sigma_{11}}}{2t}||y_2^{(2)} - X_2\beta^{(2)}||_2^2 + \frac{1}{2}\log t$$

#### S2.2 Estimating $\beta^{(i)}$ given $t$ and $\sigma_{ii}$

Numeric methods are needed for estimating $\{\beta^{(1)}, \beta^{(2)}\}$, and we use the alternating the direction method of multipliers (ADMM) [7]. ADMM introduces another constraint forcing $\beta^{(1)} = \beta^{(2)}$, which is naturally stronger than $||\beta^{(1)} - \beta^{(2)}|| < \xi$, and conveniently allows us to drop the similarity constraint of $\beta$ in Function **??** without loss in the constraint strength. With ADMM, the optimization involves solving the following three functions iteratively:

$$\begin{aligned}\beta^{(1)} =_{\beta^{(1)}} & \frac{\widehat{\sigma_{22}}}{2t}||y_1^{(1)} - X_1\beta^{(1)}||_2^2 + \lambda_1||\beta^{(1)}||_1^1 \\ & + \rho||\beta^{(1)} - \beta^{(2)}||_2^2 + \Lambda^T(\beta^{(1)} - \beta^{(2)})\end{aligned} \tag{1}$$

$$\begin{aligned}\beta^{(2)} =_{\beta^{(2)}} & \frac{\widehat{\sigma_{11}}}{2t}||y_2^{(2)} - X_2\beta^{(2)}||_2^2 + \lambda_2|\beta^{(2)}||_1^1 \\ & + \rho||\beta^{(1)} - \beta^{(2)}||_2^2 + \Lambda^T(\beta^{(1)} - \beta^{(2)})\end{aligned} \tag{2}$$

$$\Lambda = \Lambda + \rho(\beta^{(1)} - \beta^{(2)}) \tag{3}$$

where $\rho$ is another parameter introduced by ADMM. Function 1 and Function 2 can be solved via proximal gradient descent [8] where the $\ell_1$ regularization term is regarded as the proximal operator.

### S3 Convergence Proof

We proceed to offer some convergence guarantee of our iterative updating method. For the two-stage alternative optimization algorithm described in the algorithm section, we develop a reasonable proof for the convergence when modeling the objective function satisfying some weakly constrained conditions and viewing the outer iterations as block coordinate descent method for nondifferentiable minimization [9].

First, we reinterpret the main cost function as an object function over a spanned vector space composed of vectors in the form: $(\beta^{(1)};\beta^{(2)};t)$, which is generated by concatenating $\beta^{(1)},\beta^{(2)},t$ together. Therefore, the variables in the optimization problem are adapted to a block coordinate framework as the following special form:

$$f(\beta^{(1)},\beta^{(2)},t) = f_0(\beta^{(1)},\beta^{(2)},t) + \sum_{\beta^{(k)}\in\beta^{(1)},\beta^{(2)}} f_k(\beta^{(k)})$$

where

$$\begin{aligned} f_0(\beta^{(1)},\beta^{(2)},t) =& \frac{\widehat{\sigma_{22}}}{2t}||y_1^{(1)} - X_1\beta^{(1)}||_2^2 \\ &+ \frac{\widehat{\sigma_{11}}}{2t}||y_2^{(2)} - X_2\beta^{(2)}||_2^2 \\ &+ \frac{1}{2}\log t + \rho||\beta^{(1)} - \beta^{(2)}||_2^2 \\ f_k(\beta^{(k)}) =& \lambda_k||\beta^{(k)}||_1^1 \\ \sum_{\beta^{(k)}\in\beta^{(1)},\beta^{(2)}} f_k(\beta^{(k)}) =& \lambda_1||\beta^{(1)}||_1^1 + \lambda_2||\beta^{(2)}||_1^1 \end{aligned}$$

In this form, we will declare that nondifferentiable part of $f$ is separable, *i.e.*, $f_k$ only depends on one block coordinate $\beta^{(1)}$ or $\beta^{(2)}$ separately. We refer to each $\beta^{(1)},\beta^{(2)},t$ as a coordinate block of $x = (\beta^{(1)},\beta^{(2)},t) = (\beta^{(1)}_{(1)},\beta^{(1)}_{(2)},...,\beta^{(1)}_{(n)},\beta^{(2)}_{(1)},...\beta^{(2)}_{(n)},t)$, where $\beta^{(k)}_{(i)}$ is a naïve scalar for the Euclidean coordinates $(k = 1,2)$, $n$ is the dimension illustrated in Section ??. Next, we will show that each cluster point of the iterates generated by the (block) coordinate descent method is a stationary point of $f$, provided that $f_0$ has a certain convex or continuous properties as below.

For deriving sufficient conditions, the following definitions are required:

- For any $h : R^m \to R$, we use dom $h$ to denote the effective domain of $h$:

$$\text{dom } h = \{x \in R^m | h(x) \leqslant \infty\}$$

- For any $x \in$ dom $h$ and any $d \in R^m$, we denote the (lower) directional derivative of $h$ at $x$ in the direction $d$ by

$$h'(x;d) = \liminf_{\lambda\downarrow 0}[h(x+\lambda d) - h(x)]/\lambda$$

- $h$ is quasiconvex if

$$h(x+\lambda d) \leqslant \max\{h(x), h(x+d)\}$$

  for all $x, d$ and $\lambda \in [0,1]$;

- $h$ is hemivariate if h is not constant on any line segment belonging to *domh*.
- $h$ is lower semicontinuous at a point $x_0$, if the function values for arguments near $x_0$ are either close to $h(x_0)$ or greater than $h(x_0)$.

Then we are able to check properties of our objective functions:

- $f_0$ is continuous on $domf_0$
- For each $x \in \{\beta, t\}$, function $x \to f(\beta, t)$ is quasiconvex and hemivariate.
- $f_0, f_1, f_2$ are lower semicontinuous on its effective domain.

According to the Proposition in [9], when these conditions are met, we can prove the convergence effectiveness of our algorithm. Also due to only two block coordinates, we performed an efficient and accurate iterative optimization procedure for the estimation of $\beta^{(1)}$, $\beta^{(2)}$, and $t$.

### S4 Instructions of Using the Software CMM

Download and following the installation instruction at https://github.com/HaohanWang/CMM. One can also use CMM software as a stand alone script without installation.
Run

```
python cmm.py --help
```

for usage instructions as following:

```
Options:
-h, --help            show this help message and exit

Data Options:
  --file1 INPUT FILE ONE        choices of input file type
  --file2 INPUT FILE TWO        name of the input file

Model Options:
  --lambda=LMBD         the weight of the penalizer. If neither lambda or snum
                        is given, cross validation will be run.
  --snum=SNUM           the number of targeted variables the model selects. If
                        neither lambda or snum is given, cross validation will
                        be run.
  -s                    Stability selection
  -q                    Run in quiet mode
  -m                    Run without missing genotype imputation
```

**Recommended Usage:**

```
python cmm.py --file1 ./data/mice1.plink --file2 ./data/mice2.plink -m --snum 20
```

This command will run coupled mixed model and identify around 20 shared SNPs that are associated with each phenotype and jointly responsible for both phenotypes. The results are stored in data/mice1.plink.out and data/mice2.plink.out respectively

### S5 Additional Simulation Experiment Information

#### S5.1 Data Generation

We use SimPop [10] to simulate the SNPs of the two independently collected data sets. For each data set, we simulate 10000 SNPs for 500 samples from five different populations with migration behaviors. Each population also unevenly splits into five sub-populations. Therefore, it can be seen as these 500 samples are from 25 regions (denoted as $G$) out of five continents. No linkage disequilibrium is introduced explicitly during the simulation. These two SNP data sets are denoted as $X_1$ and $X_2$, following the notation described in Section **??**. $X_1$ and $X_2$ are both $500 \times 10000$ matrices.

Then we generate two sparse vectors (i.e. $\beta^{(1)}$ and $\beta^{(2)}$) as effect sizes of the association between SNPs ($X_1$, $X_2$, respectively) and phenotypes which are sampled from a Gaussian distribution $N(0, 25)$. The sparsity of these two vectors are determined by the percentage of coefficients that are non-zeros, varying in $\{0.1\%, 0.5\%, 1\%\}$. Additionally, we make sure that these two sparse vectors have an overlap of support (*i.e.,* SNPs with non-zero coefficients). The percentages of the overlapping supports range in $\{0.25, 0.5, 0.75, 0.95\}$. Hence, there are 12 sets of simulation experiments for each data set.

To introduce confounders such as population stratification and family structure, we use the aforementioned regions $G$ to affect the phenotypes differently (the effects of these regions are denoted as $\gamma$, sampled from a Gaussian distribution $N(0, 100)$). To simulate data collection confounders, we introduce a scalar $e$, sampled from a Gaussian distribution $N(0, 1)$. Therefore, the overall signal-to-noise ratio of $\beta$ is $25/(100 + 1)$, which is roughly 0.25.

Finally, we have the responses as:

$$r_1^{(1)} = X_1\beta^{(1)} + G_1\gamma_1 + e_1$$
$$r_2^{(2)} = X_2\beta^{(2)} + G_2\gamma_2 + e_2$$

Then, we transform these continuous phenotypes into binary phenotypes via Bernoulli sampling with the outcome of the inverse logit function $(g^{-1}(\cdot))$ over current responses. Therefore, we have:

$$y_1^{(1)} = \mathrm{Ber}(g^{-1}(r_1^{(1)}))$$
$$y_2^{(2)} = \mathrm{Ber}(g^{-1}(r_2^{(2)}))$$

We run the experiments with 10 different random seeds.

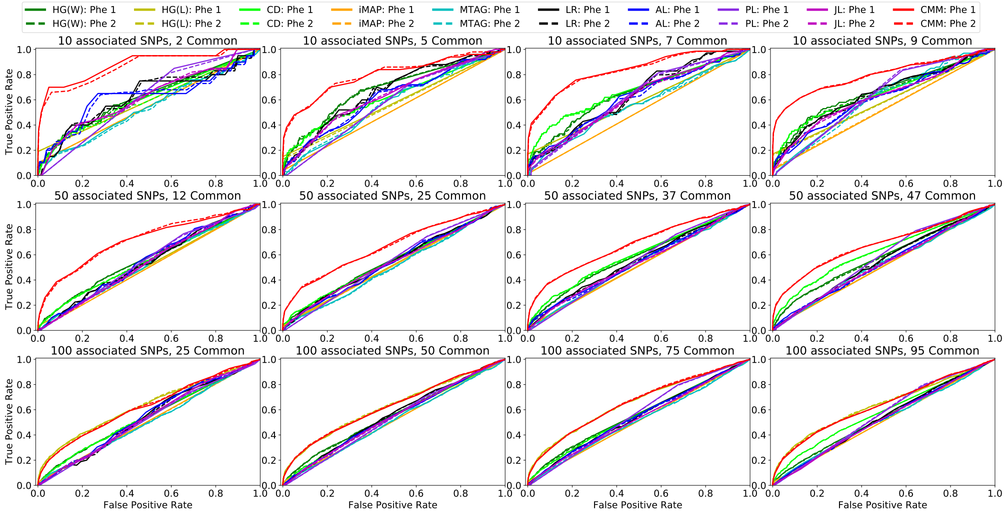


Figure S1: Roc curves of comparing methods in terms of identifying the SNPs that are responsible for each phenotype.

#### S5.2 ROC Curves for Identifying SNPs Separetely

Furthermore, we plot the results of the ROC curves of the compared methods in term of their ability to find the associated SNPs separately for each data set, which are shown in Fig. S1, where “Phe 1” and “Phe 2” stands for the two phenotypes, respectively. Notably, it is not surprising to see that the two curves for the two phenotypes almost overlap with each other. This happens because i) the two data sets were generated by using the same data generation protocol with two different sets of parameters, and ii) the results reported are averaged results across 10 different random seeds. Moreover, we notice that, these curves exhibit the same patterns as those in Fig. **??**, which is also not surprising for most competing methods when the overlapping set is only selected as the intersection of the support of the two estimated coefficients.

The results shown in these curves, along with those in Fig. **??**, indicate that CMM can identify the overlapping SNPs while also maintaining strong performance when finding associated SNPs from the two data sets independently. In this figure, we only plot the overlapping SNPs CD identifies, since CD is unable to identify any SNPs for the two phenotypes separately.

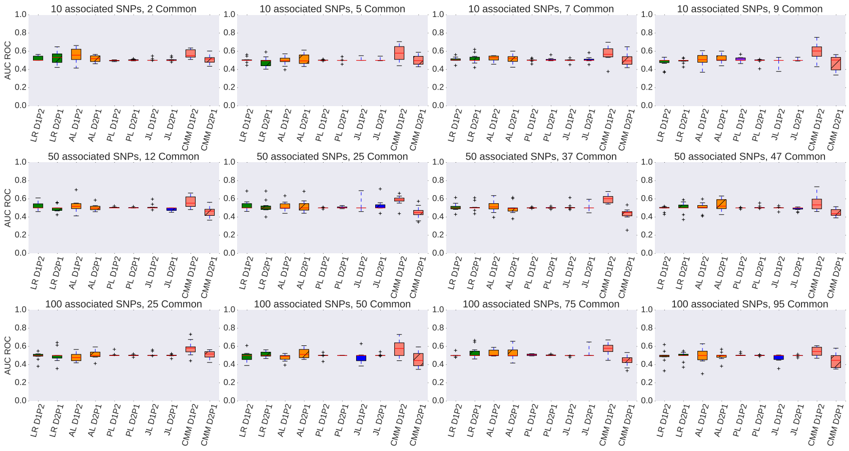


Figure S2: Area under ROC curve of cross data set phenotype prediction. "D$i$P$j$" denotes the prediction preformance of Phenotype $j$ in Data Set $i$.

#### S5.3 Prediction Performance

In addition to the performance in identifying the associated SNPs, we also test how these identified associated SNPs can help in predicting the phenotype that is not seen by the model. In other words, after we estimate the $\widehat{\beta^{(1)}}$ and $\widehat{\beta^{(2)}}$ with $X_1$, $X_2$, $y_1^{(1)}$, and $y_2^{(2)}$, we continue to test how $\widehat{\beta^{(2)}}$ helps in prediction $y_1^{(2)}$ (denoted as D1P2) and how $\widehat{\beta^{(2)}}$ helps in prediction $y_2^{(1)}$ (denoted as D2P1). Because the testing phenotype may not be balanced in terms of the case/control samples, we evaluate the performance with area under ROC curve. We only evaluate the multivariate regularized regression methods because univariate testing methods are not designed for prediction. We test the average performance of same set of hyperparameter choices as the association study experiment. The results of 10 random seeds are visualized via standard box plot in Fig. S2.

As Fig. S2 shows, due to the existence of confounding factors, prediction across data set is an extremely difficult task. Most methods only behave marginally better than random chances. CMM shows an advantage over the competing methods as it shows a higher median performance and also higher third-quartile/maximum performances in most cases. Interestingly, Adaptative Lasso also behaves resonabley well.

### S6 Additional Literature Support of Real Data Analysis

*rs2131691* which resides in the *ANO3* gene is identified by our method as the SNP associated with both AD and SUD with the most significant effect sizes. It is widely known that in the brain of AD patients, the metabolism of amyloid-$\beta$ (A$\beta$) and tau proteins is dysregulated and crucially influenced by autophagy, thus autophagy plays an important role in AD [11]. *ANO3* is a gene that encodes the protein anoctamin 3, which belongs to a family of calcium-activated chloride channels [12]. Although no clear evidence showing the role of calcium-activated chloride channels in autophagy, other chloride channels including chloride voltage-gated channels, cystic fibrosis transmembrane conductance regulator and volume-regulated anion channel have been proposed to regulate autophagy [13]. Therefore, it is worth experimentally exploring the potential role of *ANO3* in autophagy and AD. Besides that, *ANO3* is reported to be associated with AD *in silico* [14]. To the best of our knowledge, there is no previous literature that associates *ANO3* with SUD. However, evidence has shown that mutations in *ANO3* produced functional changes that affected striatal signal transduction pathways [15, 16] and also that dysfunction of striatal signal transduction pathways is implicated in SUD [17]. These observations suggest that *rs2131691* is connected to SUD.

*rs1709317*, which resides in *KLHL29*, is identified by our method as the 8$^{\text{th}}$ SNP and 5$^{\text{th}}$ SNP associated with AD and SUD respectively. We do not find strong previous evidence that supports the association between *KLHL29* and either AD or SUD. However, as shown in Table 2 (main manuscript), *rs1709317* is significantly different in the case vs. control samples in both AD and SUD.

*rs4713797*, which resides in *PACSIN1*, is identified as the 10$^{\text{th}}$ SNP and 6$^{\text{th}}$ SNP associated with AD and SUD, respectively. *PACSIN1* encodes protein kinase C and casein kinase substrate in neurons protein 1, which plays a role in synaptic neurotransmission and neuroplasticity. Specifically, *PACSIN1* is demonstrated to regulate the dynamics of AMPA receptor trafficking [18] and involved in the reorganization of the actin cytoskeleton and neuron morphogenesis [19]. Neurotransmission and neuroplasticity play essential roles in drug addiction process [20, 21], which suggests the potential role of *PACSIN1* in SUD. A study investigated whether DNA methylation patterns in early life are prospectively associated with SUD in adolescence [22] also demonstrated the association between *PACSIN1* and SUD. In addition, *PACSIN1* is reported to be an important Tau binding partner in regulating microtubule dynamics and forming axonal plasticity [23], and Tau has been correlated with AD [24]. Besides that, *PACSIN1* has been implicated in various neurodegenerative disorders, including Huntington and AD [25, 26, 27].

*rs1057744*, which resides in *JAG2*, is identified by the CMM method as the 11$^{\text{th}}$ and 16$^{\text{th}}$ SNP associated with AD and SUD, respectively. It also shows significant difference between the case vs. control samples in both AD and SUD (Table 2, main manuscript). *JAG2* has been reported to interact with NOTCH2 [28], and is involved in Notch signaling pathway according to KEGG database, several studies have demonstrated the connection of Notch signaling and AD [29, 30, 31]. [32] reported the association of *JAG2* with late-onset AD. For SUD, a study has shown that the Notch signaling pathway is closely related to the abuse of opioids, explains the alterations triggered in the early stages of drug addiction [33].

### S7 Additional Real Data Results

#### S7.1 Substance use disorder

Table S1: **The top SNPs that our method identifies in substance use disorder study with joint study in AD** The SNPs are ranked by the absolute values of their estimated effect sizes. SNPs in bold are the ones that are shared with the results in Alzheimer's disease study. The MAFs reported in the table are calculated using the case-control substance use disorder experimental data. The information of whether a SNP is located within a region of a gene is taken from the Database for Single Nucleotide Polymorphisms (dbSNP) [34], and listed in the 'Gene' column. The literature support showed supports the association between the corresponding gene and the disease either *in vivo* or *in silico* from other independent data sets. Abbreviations: AD: Alzheimer's Disease, ALC: Alcoholism, HD: Huntington's disease, SCZ: Schizophrenia, SMK: Cigarette smoking, SUD: Substance use disorder.

| Rank | SNP | Chr | Chr. Position | MAF | Gene | Disease (Literature) |
|---|---|---|---|---|---|---|
| **1** | **rs2131691** | **11** | **26574855** | **0.47** | ***ANO3*** | **AD [14]** |
| 2 | rs11167137 | 8 | 142312847 | 0.47 | *TSNARE1* | SCZ [35] |
| 3 | rs7267819 | 20 | 53888742 | 0.48 | | |
| 4 | rs10759638 | 9 | 113280212 | 0.36 | *PRPF4* | SUD [36] |
| **5** | **rs1709317** | **2** | **23536638** | **0.36** | ***KLHL29*** | **SMK [37]** |
| **6** | **rs4713797** | **6** | **34490756** | **0.48** | ***PACSIN1*** | **AD [26], HD [25], SUD [22]** |
| 7 | rs13155209 | 5 | 10788807 | 0.38 | | |
| 8 | rs755598 | 12 | 103666012 | 0.42 | *STAB2* | SUD [38] |
| 9 | rs1997858 | 6 | 115688219 | 0.46 | | |
| 10 | rs17356935 | 7 | 106357500 | 0.44 | | |
| 11 | rs7586009 | 2 | 46871829 | 0.40 | | |
| **12** | **rs224534** | **17** | **3583408** | **0.33** | ***TRPV1*** | **AD [39], ALC [40], SUD [41]** |
| 13 | rs3750534 | 9 | 113297844 | 0.36 | *RNF183* | |
| 14 | rs2790453 | 10 | 28481089 | 0.47 | | |
| 15 | rs1415640 | 13 | 44768660 | 0.37 | | |
| **16** | **rs1057744** | **14** | **105150705** | **0.49** | ***JAG2*** | **AD [32]** |
| 17 | rs2736100 | 5 | 1286401 | 0.48 | *TERT* | |
| 18 | rs1398800 | 6 | 66487703 | 0.33 | | |
| 19 | rs2337158 | 5 | 168249731 | 0.38 | *TENM2* | |
| 20 | rs10865088 | 2 | 31304987 | 0.32 | | |

The identified SNPs for DA are listed in Table S1. There are 20 SNPs identified in total and five of these SNPs (showed in bold) are also identified to be associated with AD.

The SNP that has been identified with the strongest effect size resides within the region of gene *ANO3*. To the best of our knowledge, there is no previous literature that associates *ANO3* with DA. However, we notice evidences showing that ANO3 mutations produce functional changes that converge to affect striatal signal transduction pathways [15, 16] and [17] connected the dysfunction of striatal signal transduction pathways and DA by showing that the repeated exposure to drugs can lead to alternation of functional and structural neuroplasticity, eventually leading to addiction behavior. Chances are that mutation of *rs2131691* can lead to the consequence as the result of repeated drug exposure in terms of the dysfunction of striatal signal transduction pathways.

The $2^{nd}$ SNP resides within the region of gene *TSNARE1*, which has been reported to be associated with Schizophrenia [35], another cognitive dysfunction. The often co-occurrences between Schizophrenia and DA has raised the interest in studying the comorbidity of these two [42]. Our finding provides another evidence to the linkage between DA disorder and the cognitive dysfunctions.

The $4^{th}$ SNP resides within the region of gene *PRPF4*, which has also been reported as an association gene to substance dependence from another independently collected data set *in silico* [36].

The next SNP is identified both for DA and AD in our study. It resides within gene *KLHL29*. *KLHL29* has been reported as one of the top 25 small airway epithelium hypomethylated genes of smokers compared with nonsmokers [43], also reported to be associated with smoking cessation

[37].

The next SNP is also identified to be associated with both DA and AD and it resides within gene *PACSIN1*. The association between *PACSIN1* and substance addiction has been reported in an investigation of whether DNA methylation patterns in early life prospectively associate with substance use in adolescence conducted by [22].

The 8$^{\text{th}}$ identified SNP resides within gene *STAB2*, which has been reported to be associated with opioid dependence by [38] in another independent data set.

The 12$^{\text{th}}$ SNP is also identified to be associated with both DA and AD. It resides in the gene *TRPV1*. [41] have experimented with mice to show that *TRPV1* is related with opiate drug morphine-additive disorders. In addition, [40] showed that the deletion of *TRPV1* in mice alters behavioral effects of ethanol, which indicates the connection between *TRPV1* and Alcoholism.

The 16$^{\text{th}}$ SNP is also identified to be associated with both DA and AD in our study. The SNP resides within the region of gene *JAG2*. To the best of our knowledge, previous study did not show direct evidence about the association between *JAG2* and DA, but we notice that [44] observed significant down-regulation of *JAG2* with heavy alcohol consumption in female rhesus macaques.

#### S7.2 Alzheimer's Disease

The identified SNPs for AD are listed in Table S2 (Table is showed in the last page). There are 40 SNPs identified in total and five of these SNPs (showed in bold) are also identified to be associated with DA.

The most significantly associated SNP is the same as the most significantly associated SNP with DA and it resides within the region of gene *ANO3*, which is also reported to be associated with DA *in silico* [14].

The 2nd identified SNP resides in the gene *LRRC38*. *LRRC38* has also been reported to be associated with memory impairment, the cardinal early feature of DA, in older adults [45].

The 3rd SNP has also been reported to be associated with DA. Regarding AD, [39] has showed that the positive modulation of *TRPV1* channels can be a potential target for mitigation of AD, indicating the relationship between *TRPV1* and AD.

The 6th SNP identified resides within the region of gene *GRIP1*. Although there is no direct evidence to show that it is related to AD, [56] has shown that *GRIP1* is essential for regulation in synaptic scaling (a form of homeostatic plasticity that stabilizes neuronal firing in response to changes in synapse number and strength). Also, defects in synaptic scaling is related with AD [57]. In addition to the association with AD, we also notice evidence relating *GRIP1* with cocaine addiction [46].

The 8th SNP resides within gene *KLHL29* and it has also been identified to be associated with DA. Additionally, *KLHL29* has been reported to be associated with AD from COSMIC and DistiLD via [58][1].

The 9th, 22nd, 28th, and 35th SNPs are all in the region of gene *CSMD1*. *CSMD1* has reported to be associated with AD by [47] with copy number analysis. However, we should be cautious that they analyzed the same data set as our study, in despite of the different approaches. In addition, *CSMD1* is also evidently associated with Schizophrenia [48, 49].

The next SNP that is in the region of *PACSIN1* has also been reported to be associated with DA in our study. Additionally, *PACSIN1* has also been implicated in various neurodegenerative disorders, including Huntington and Alzheimer's diseases [25, 26, 27].

The next SNP resides in *JAG2* and it has also been identified to be associated with DA. In addition, [32] reported the association between *JAG2* with late-onset Alzheimer's disease (AD) develop psychosis.

The 14th, 26th, and 31th SNPs all reside within the region of gene *MEGF10*. [50, 51] have showed the evidence relating *MEGF10* and AD through the mechanism of amyloid$-\beta$.

The 20th resides within the region of *CPVL*, which has also identified as a candidate biomarker for AD by [52] from another independent study.

The 30th identified SNP resides in the region of *RAP52*. [53] has shown the relationship between *RAP52* and AD through the mechanism of amyloid$-\beta$.

The 32nd and 38th SNP are both in the gene *PRKG1*. Although we do not notice evidences associating *PRKG1* with AD, this gene has been implicated related disorders like Alcoholism [54] and Schizophrenia [55].

[1] https://diseases.jensenlab.org/Search

Table S2: **The top SNPs that our method identifies in Alzheimer's disease study with joint study in substance use disorder** The SNPs are ranked by the absolute values of their estimated effect sizes. SNPs in bold are the ones that are shared with the results in substance use disorder disorder study. The MAFs reported in the table are calculated using the case-control AD experimental data. The information of whether a SNP is located within a region of a gene is taken from the Database for Single Nucleotide Polymorphisms (dbSNP) [34], and listed in the 'Gene' column. The literature support showed supports the association between the corresponding gene and the disease either *in vivo* or *in silico* from other independent data sets. Abbreviations: AD: Alzheimer's Disease, ALC: Alcoholism, HD: Huntington's disease, SCZ: Schizophrenia, SMK: Cigarette smoking, SUD: Substance use disorder.

| Rank | SNP | Chr | Chr. Position | MAF | Gene | Disease (Literature) |
|---|---|---|---|---|---|---|
| **1** | **rs2131691** | **11** | **26574855** | **0.47** | ***ANO3*** | **AD [14]** |
| 2 | rs2999899 | 1 | 13502964 | 0.47 | *LRRC38* | AD [45] |
| **3** | **rs224534** | **17** | **3583408** | **0.34** | ***TRPV1*** | **AD [39], ALC [40], SUD [41]** |
| 4 | rs2944529 | 10 | 131160663 | 0.41 | *TCERG1L* | |
| 5 | rs2358126 | 2 | 11790153 | 0.47 | *LPIN1* | |
| 6 | rs975731 | 12 | 66801694 | 0.45 | *GRIP1* | SUD [46] |
| 7 | rs6063725 | 20 | 51865839 | 0.41 | | |
| **8** | **rs1709317** | **2** | **23536638** | **0.37** | ***KLHL29*** | **SMK [37]** |
| 9 | rs1700106 | 8 | 4243970 | 0.48 | CSMD1 | AD [47], SCZ [48, 49] |
| **10** | **rs4713797** | **6** | **34490756** | **0.48** | ***PACSIN1*** | **AD [26], HD [25], SUD [22]** |
| **11** | **rs1057744** | **14** | **105150705** | **0.50** | ***JAG2*** | **AD [32]** |
| 12 | rs1223552 | 6 | 13173601 | 0.49 | *PHACTR1* | |
| 13 | rs473015 | 10 | 17942519 | 0.45 | | |
| 14 | rs2546078 | 5 | 127332991 | 0.49 | *MEGF10* | AD [50, 51] |
| 15 | rs6088244 | 20 | 33522398 | 0.48 | *CBFA2T2* | |
| 16 | rs12611086 | 19 | 34136027 | 0.43 | | |
| 17 | rs4766335 | 12 | 5059683 | 0.46 | | |
| 18 | rs11134717 | 5 | 171698478 | 0.46 | | |
| 19 | rs7308687 | 12 | 5249169 | 0.50 | | |
| 20 | rs317726 | 7 | 29081595 | 0.48 | *CPVL* | AD [52] |
| 21 | rs2730753 | 12 | 118734169 | 0.46 | | |
| 22 | rs1481720 | 8 | 3756931 | 0.33 | *CSMD1* | AD [47], SCZ [48, 49] |
| 23 | rs2155440 | 11 | 108559049 | 0.50 | *EXPH5* | |
| 24 | rs10418480 | 19 | 57243687 | 0.49 | *ZNF805* | |
| 25 | rs4604527 | 9 | 82909654 | 0.39 | | |
| 26 | rs10065046 | 5 | 127233347 | 0.43 | *MEGF10* | AD [50, 51] |
| 27 | rs910397 | 20 | 33707735 | 0.48 | *PXMP4* | |
| 28 | rs12543111 | 8 | 3756707 | 0.26 | *CSMD1* | AD [47], SCZ [48, 49] |
| 29 | rs11126690 | 2 | 78971494 | 0.36 | | |
| 30 | rs10744729 | 12 | 915428 | 0.42 | *RAD52* | AD [53] |
| 31 | rs27652 | 5 | 127348191 | 0.49 | *MEGF10* | AD [50, 51] |
| 32 | rs4568954 | 10 | 51750701 | 0.48 | *PRKG1* | ALC [54], SCZ [55] |
| 33 | rs4606813 | 18 | 76766320 | 0.45 | | |
| 34 | rs9878318 | 3 | 190296916 | 0.45 | | |
| 35 | rs4875786 | 8 | 3779552 | 0.37 | *CSMD1* | AD [47], SCZ [48, 49] |
| 36 | rs714180 | 7 | 116678948 | 0.47 | *MET* | |
| 37 | rs4743713 | 9 | 96307662 | 0.44 | | |
| 38 | rs4466778 | 10 | 51747293 | 0.42 | *PRKG1* | ALC [54], SCZ [55] |
| 39 | rs7513520 | 1 | 62823261 | 0.44 | *ATG4C* | |
| 40 | rs4238945 | 16 | 26963686 | 0.45 | | |
